# Analysis of Metabolic Stability under Environmental Perturbations Using a Kinetic Model

**DOI:** 10.1101/2025.08.27.672763

**Authors:** Atsuki Hishida, Yusuke Himeoka, Chikara Furusawa

**Affiliations:** Graduate School of Science, Kyoto University, Kyoto, Japan; Universal Biology Institute, The University of Tokyo, Tokyo, Japan; RIKEN Center for Biosystems Dynamics Research, Kobe, Japan

## Abstract

Cells robustly maintain metabolic function despite environmental fluctuations affecting reaction kinetics, yet kinetic models often exhibit fragility. We investigated this discrepancy by examining the effects of temperature changes on a kinetic model of *Escherichia coli* central metabolism. A gradual temperature decrease caused an abrupt transition to a state with sharply reduced ATP production efficiency, triggered by an elevated ATP/ADP ratio creating a glycolysis bottleneck. Introducing a rapid ATP-ADP exchange reaction prevented this ATP/ADP ratio surge and sustained high ATP production efficiency across temperatures. Furthermore, altering enzyme abundances similarly maintained high ATP production, with predicted enzyme regulation aligning with experimental observations in *E. coli* at low temperatures. Our findings indicate that balancing key cofactors, such as the ATP/ADP ratio, is crucial for preserving metabolic stability under environmental perturbations.

## Introduction

Cellular metabolism comprises thousands of reactions that together generate energy, build biomolecules, and recycle cellular components. As external conditions fluctuate, cells adjust their metabolic state to maintain function. For example, bacteria switch between fermentative and respiratory regimes, reroute carbon through the pentose phosphate pathway (PPP) under oxidative stress, and upregulate anaplerotic reactions when nutrients are scarce (Christodoulou et al., 2018; Sawers, 2025; Sudarsan et al., 2025). This plasticity arises from regulatory mechanisms such as enzyme reallocation and allosteric modulation, enabling cells to occupy distinct, environment-specific metabolic states while sustaining growth (Gruber et al., 2025; Miyakoshi, 2024).

Mathematical models of metabolic dynamics have been widely used to probe this plasticity (El-Samad et al., 2005; Stephanopoulos et al., 1998; Strutz et al., 2019). Such models allow analysis of how steady states shift as parameters vary. A recurring finding is that steady-state stability is often fragile to parameter changes. For instance, a kinetic model of *E. coli* core metabolism was stable for only a small fraction of flux and concentration sets consistent with its kinetic parameters (Chakrabarti et al., 2013). Similarly, Steuer et al. (2006) sampled thousands of parameter sets for a kinetic model of yeast glycolysis and found that over 80% of resulting steady states were unstable. These narrow stability ranges suggest that environmental perturbations can readily destabilize cellular metabolism. Yet, despite the fragility predicted by models, real cells maintain stable metabolic states as temperature and nutrient availability change (Phadtare, 2004; Phadtare & Inouye, 2008; Riccardi et al., 2023; Zhang & Gross, 2021). This discrepancy implies the existence of mechanisms that stabilize metabolism. Indeed, cells adjust enzyme levels and pathway activities in response to environmental cues (Jozefczuk et al., 2010; Schmidt et al., 2016; Ye et al., 2012; Zhao et al., 2024). However, how such regulation preserves dynamic stability remains unclear.

Among environmental perturbations, temperature shifts directly and broadly affect metabolism (Noll et al., 2020; Wendering & Nikoloski, 2023). Temperature modulates the rate constants of all intracellular reactions, with magnitudes set by each reaction’s activation free energy (Evans & Polanyi, 1935; Eyring, 1935). Because forward and reverse reactions need not respond identically, temperature changes can alter both rates and reversibility, reshaping flux throughout the network. Such nonuniform effects may destabilize steady states. Clarifying how cells prevent this outcome would reveal general principles of metabolic robustness.

Here, we investigate mechanisms that keep complex metabolic systems stable under temperature-induced parameter perturbations. Using a kinetic model of *E. coli* central metabolism constructed from experimental data, we varied model parameters continuously with temperature and observed a bifurcation: the system transitioned discontinuously to a new steady state. The post-bifurcation state exhibited an elevated ATP/ADP ratio and reduced ATP production efficiency. Imposing operations that maintain the ATP/ADP ratio during the temperature shift eliminated the bifurcation and preserved ATP production efficiency even at low temperatures. Moreover, adjusting enzyme abundances to increase ATP-utilizing reaction rates stabilized the ATP/ADP ratio. These results indicate that maintaining the balance of key cofactors is critical for stabilizing metabolic systems under parameter perturbations caused by environmental change.

## Results

### Temperature decrease induces a bifurcation

To examine how a low-temperature shift affects the metabolic state of *E. coli* central metabolism, we simulated metabolic dynamics using rate constants adjusted for different temperatures. Figure 1 shows the reaction network of the kinetic model used in this study (see Materials and Methods for details). Rate constants at each temperature were calculated using the Eyring–Polanyi equation. We focused on temperature downshifts because heating can cause irreversible enzyme inactivation, which is not captured by the standard Eyring–Polanyi formulation (Arcus & Mulholland, 2020; Noll et al., 2020). Activation free energies (Δ*G*^‡^) for individual reactions were obtained from parameter values fitted to the *in vivo* metabolic state at 37 °C reported by Khodayari et al. (2014) (see Materials and Methods for the details). Without a temperature shift, the system converges to a stable steady state in which glucose serves as the sole carbon source and ATP is produced via glycolysis, the TCA cycle, and the electron transport chain (ETC). We refer to this as the “original steady state” throughout the paper.

**Figure 1.**
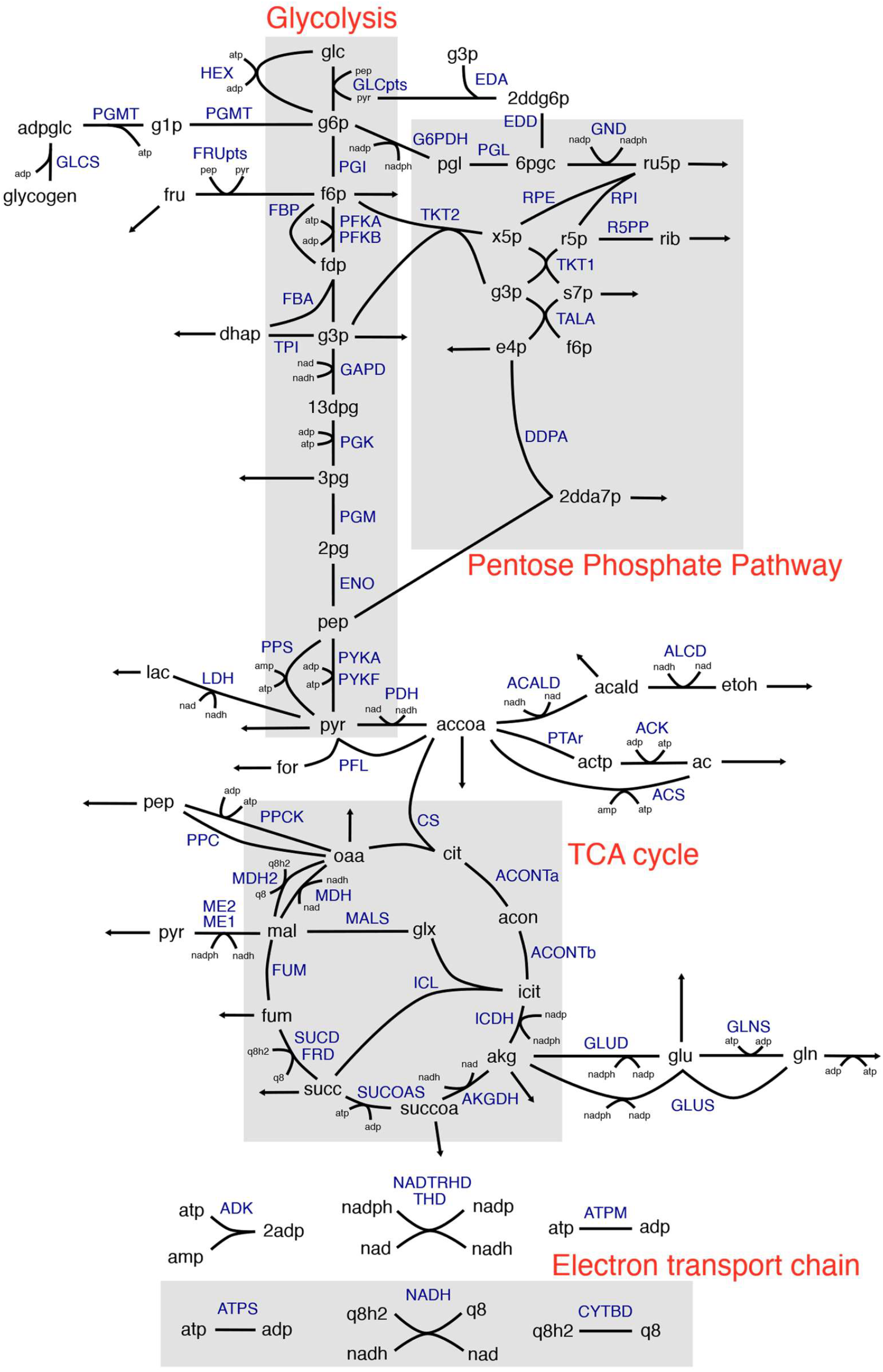
Reaction network of the kinetic model used in our simulations. Although each reaction was decomposed into elementary steps for the simulations, the network is shown here with the pre-decomposition reactions for clarity.

When the temperature is lowered from 37 °C to 0 °C, each rate constant decreases according to its activation free energy. As a result, the distribution of rate constants shifts, and the median value decreases by about 10-fold (Figure 2A). We simulated system dynamics using the rate constants at 0 °C, starting from the original steady state. Numerical simulations showed that metabolite concentrations converged to a new stable steady state (Figure 2B). Compared with the original state, metabolite concentrations in this new steady state changed by factors ranging from 10^−9^ to 10^4^— far greater than the fold change observed in the median rate constant.

**Figure 2.**
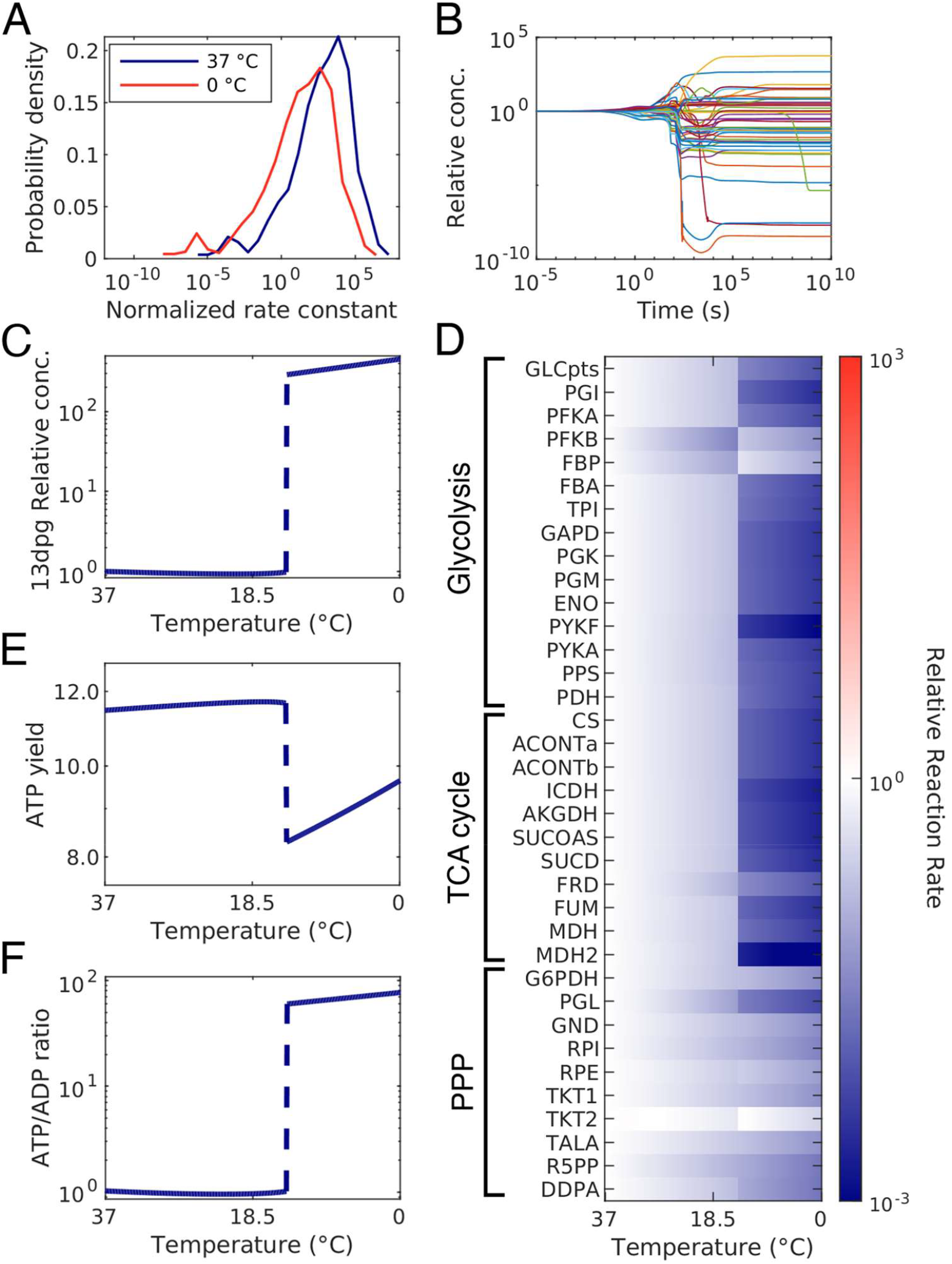
(A) Distribution of reaction rate constants in the system. (B) Time courses of relative metabolite concentrations with all rate constants fixed at their values for 0 °C and initial concentrations set to the original steady state; each line represents an individual metabolite. (C) Steady-state concentration of D-glycerate 1,3-bisphosphate at each temperature. (D) Relative reaction rates at steady state for each temperature, normalized to the rates at the original steady state. (E, F) Steady-state ATP yield (E) and ATP/ADP ratio (F) at each temperature.

To determine the temperature at which this substantial shift occurs, we tracked the steady state as the temperature was gradually decreased from 37 °C to 0 °C. At each step, system dynamics were integrated up to *t* = 10^10^ *s* to ensure convergence, and the resulting concentrations were used as initial conditions for the next step, with temperature reduced by 0.1 °C. At 14.2 °C, the steady state shifted discontinuously, with abrupt changes in the concentrations of all metabolites (Figure 2C shows one example; Supplementary Figure S1 presents all metabolite concentrations at the converged state across temperatures). Reaction rates also changed discontinuously, and relative changes compared with the original steady state are shown in Figure 2D.

This discontinuous shift could arise from either the disappearance or destabilization of the original steady state, or from a switch between coexisting attractors. To discriminate between these possibilities, we enumerated multiple stable steady states across temperatures including the transition point. We generated 10,000 random initial conditions by scaling each chemical concentration of the original steady state by 10^*U*(−3.0,3.0)^. Here *U*(*a, b*) denotes a uniform distribution ranging from *a* to *b*. We sampled initial conditions uniformly in the logarithmic scale over the interval (10^−3^, 10^3^). See Materials and Methods for details of sampling procedure. Starting from each of these points, we numerically integrated the system dynamics to *t* = 10^10^ *s* and checked convergence for all cases.

This procedure revealed three coexisting stable steady states above 14.2 °C and two below it. These results indicate that the discontinuous shift arises from a bifurcation, in which the physiologically relevant steady state above 14.2 °C becomes destabilized as temperature decreases. Bifurcation diagrams for all small molecules are provided in Supplementary Figure S1. Notably, the disappearing steady state corresponds to the physiological *E. coli* state, characterized by aerobic respiration and ATP production from glucose.

Examination of the steady state after the bifurcation revealed substantial changes in metabolic fluxes and ATP production efficiency. The discontinuous shift caused glycolytic and TCA cycle fluxes to decrease by factors of 100–1,000, while flux through the pentose phosphate pathway (PPP) increased (Figure 2D). As a result, ATP yield—defined as the sum of ATP-producing fluxes divided by the glucose uptake rate—declined following the temperature-induced bifurcation (Figure 2E).

To investigate the cause of the decreased ATP yield, we examined metabolite concentrations along glycolysis and the TCA cycle. While most metabolite levels decreased after the temperature-induced bifurcation, the concentration of D-glycerate 1,3-diphosphate (13dpg) increased by two orders of magnitude compared with its pre-bifurcation level (Figure 2C; Supplementary Figure S1). Despite this elevation, the reaction degrading 13dpg—catalyzed by phosphoglycerate kinase (PGK), which converts 13dpg and ADP to 3-phosphoglycerate and ATP in glycolysis (Figure 1)—showed a reduced rate. This reduction was caused by a ~40-fold increase in the ATP/ADP ratio, driven by elevated ATP and decreased ADP concentrations (Figure 2F; Supplementary Figure S1). Because the PGK reaction lies near the center of glycolysis, its slowdown created a bottleneck that diminished flux through both glycolysis and the downstream TCA cycle.

### Fixing the ATP/ADP ratio prevents the temperature-induced bifurcation

The above results suggested that a sudden rise in the ATP/ADP ratio can trigger the temperature-induced bifurcation. To investigate this relationship, we maintained the ATP/ADP ratio nearly constant by introducing a rapid, reversible ATP–ADP exchange reaction into the system. Because the ATP/ADP ratio at the steady state before the bifurcation was approximately 1:1, the forward and reverse rate constants were set to 10^6^, far exceeding the median values of other reactions in the model. A detailed procedure for fixing the ATP/ADP ratio during temperature decrease is provided in Materials and Methods.

The modified model with the ATP–ADP exchange reaction maintained a single, stable steady state under the 37 °C parameter conditions. The resulting concentrations of 13dpg, the ATP/ADP ratio, and the ATP yield were similar to those of the original steady state (Figure 3A–C). With the exchange reaction, lowering the temperature from 37 °C to 0 °C no longer caused a sharp rise in 13dpg concentration or discontinuous changes in pathway fluxes, as shown by the relative changes in reaction rates (Figure 3D). Consistently, the ATP/ADP ratio remained near unity, and the ATP yield showed no abrupt decline across the temperature range (Figure 3B, C). To test for multiple attractors at each temperature, we generated 10,000 random initial points by scaling each metabolite concentration of the original steady state by 10^*U*(−3.0,3.0)^ and simulated the dynamics until convergence. All trajectories converged to a single steady state at every temperature. These results indicate that, if the ATP/ADP ratio is fixed at approximately 1.0, the system does not undergo bifurcation, even though individual reaction rate constants still vary with temperature.

**Figure 3.**
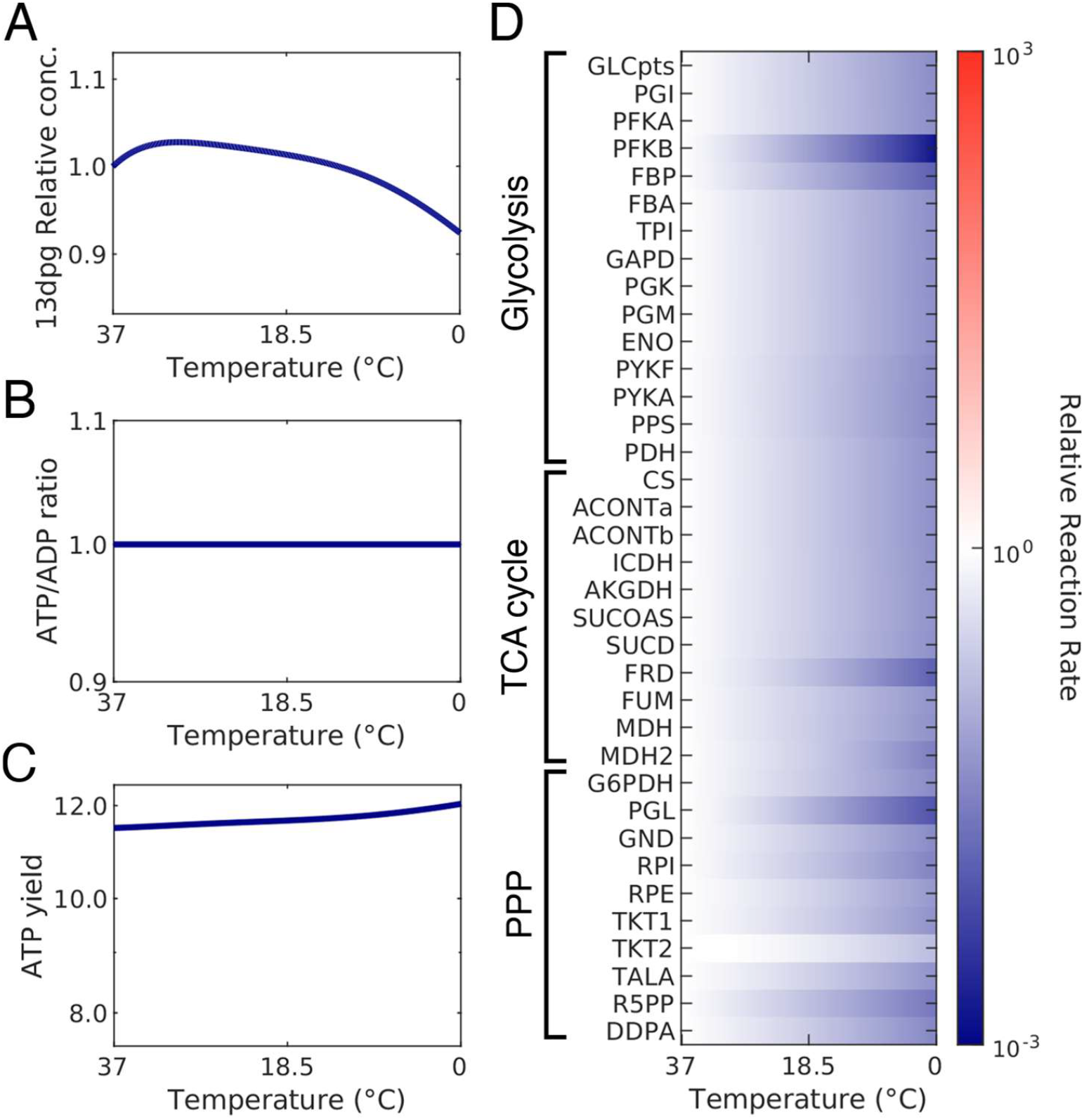
Changes in steady states of the modified model with ATP/ADP ratio homeostasis. (A) Steady-state concentration of D-glycerate 1,3-bisphosphate at each temperature. (B, C) Steady-state ATP/ADP ratio (B) and ATP yield (C) at each temperature. (D) Relative changes in reaction rates at steady state for each temperature.

### Maintaining the ATP yield by adjusting enzyme abundances

The analysis in the previous section demonstrated that the ATP yield can be maintained at a high level by fixing the ATP/ADP ratio using a rapid, reversible ATP–ADP exchange reaction. However, cells do not naturally possess such a reaction, and precise control of the ATP/ADP ratio in this manner is biologically implausible. Instead, cells adjust intracellular enzyme abundances to adapt to diverse environmental perturbations, including temperature fluctuations, osmotic stress, oxidative conditions, and nutrient limitations (Jozefczuk et al., 2010; Schmidt et al., 2016; Ye et al., 2012; Zhao et al., 2024). We therefore investigated whether ATP yield could be maintained under decreasing temperatures by modulating enzyme abundances. To this end, we sampled steady states with varied enzyme levels under low-temperature conditions and identified enzyme abundances that supported a high ATP yield.

A naïve approach of randomly varying enzyme levels and integrating the system dynamics until convergence for each set would require computationally intensive simulations. To overcome this, we developed a faster method to sample numerous low-temperature steady states. Because every reaction in our model is decomposed into elementary steps, each reaction rate function depends on at most one or two variables and follows mass-action kinetics. Consequently, the rate functions are at most quadratic polynomials, with the second variable, if present, corresponding to the concentration of a small-molecule substrate. By fixing the concentrations of small molecules near their values in the original steady state, the rate functions become linear, transforming the system of differential equations into a linear one. Steady states of this linear system can then be obtained simply by solving a linear equation (equation (7) in Materials and Methods).

The linearized system has infinitely many steady states. To sample one, we first selected a random reference point of enzyme abundances *x*_*ref*_, near those of the original steady state. Each concentration in the original steady state was multiplied by a value drawn from a log-normal distribution over the interval (1/3, 3.0). Broadening this range reduces the fraction of sampled states that are stable, so we chose the widest interval that still yielded stable solutions with reasonable frequency. We then solved a linear optimization problem to identify the steady state closest to *x*_*ref*_, subject to additional constraints: (1) the glucose transport reaction must proceed inward while other transport reactions act outward, and (2) the total enzyme amount must equal that of the original steady state. These transport constraints were imposed to ensure the biological plausibility of the sampled steady states. A mathematical formulation and detailed description of the sampling procedure are provided in Materials and Methods. This approach samples steady states more efficiently than the naïve method. Because stability was not enforced during sampling, we assessed it post hoc using the Jacobian matrix at each steady state. Materials and Methods also describe how steady states were classified as stable based on the Jacobian.

Using this method, we sampled 5,000 steady states with rate constants corresponding to 0 °C and visualized the 94-dimensional enzyme profiles in two dimensions via principal component analysis (PCA; Figure 4A). Among all computed solutions, the stable steady states were concentrated in a specific region, where TCA cycle enzyme levels were elevated. Figure 4B shows the sum of TCA enzyme abundances at each steady state in the PCA plot. These results suggest that increasing TCA cycle enzyme concentrations is important for stabilizing the metabolic state under low-temperature conditions.

**Figure 4.**
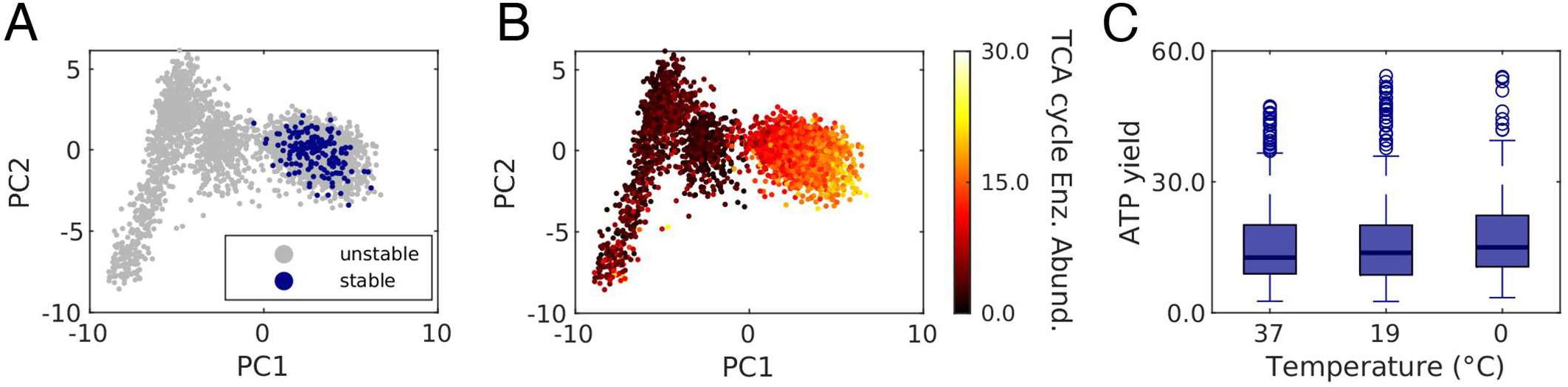
(A) PCA plot of 94-dimensional enzyme abundances at steady states obtained with parameters for 0 °C. Explained variance: PC1, 16.9%; PC2, 4.7%. Stability of each steady state is indicated by color: blue, stable; grey, unstable. (B) TCA cycle enzyme abundances (color-coded) at steady states under low-temperature conditions. (C) Box plot of ATP yield for stable steady states at each temperature.

The above results demonstrate that adjusting enzyme abundances allows the system to achieve stable steady states near the original steady state, even under low-temperature conditions. However, ATP yields at these stable states are not necessarily high. To evaluate ATP yield under low temperatures, we sampled stable steady states across temperatures from 37 °C to 0 °C using the same method and assessed their ATP production. The ATP yield remained high even at low temperatures. Figure 4C shows box plots of ATP yields for stable steady states at 37 °C, 19 °C, and 0 °C. These findings indicate that cells can maintain a high ATP yield during a low-temperature shift by modulating enzyme abundances.

In practice, cells may not experience a sudden drop in temperature but rather undergo gradual cooling. We therefore investigated which enzyme reallocations would allow the system to maintain a high ATP yield as temperature decreases from 37 °C to 0 °C. To do this, we fixed the target concentrations of free enzymes and substrate-enzyme complexes (*x*_*ref*_), and computed the steady-state enzyme abundances at each temperature. Along this path, enzyme levels at stable steady states changed with temperature: TCA cycle enzymes increased, whereas ETC enzyme levels decreased. Figures 5A and 5B show the summed abundances of TCA cycle and ETC enzymes at stable steady states across temperatures. This trend is consistent with experimental observations that cold-adapted cells upregulate TCA cycle enzymes and downregulate ATP synthase subunits (Knapp et al., 2024; Phadtare & Inouye, 2004).

**Figure 5.**
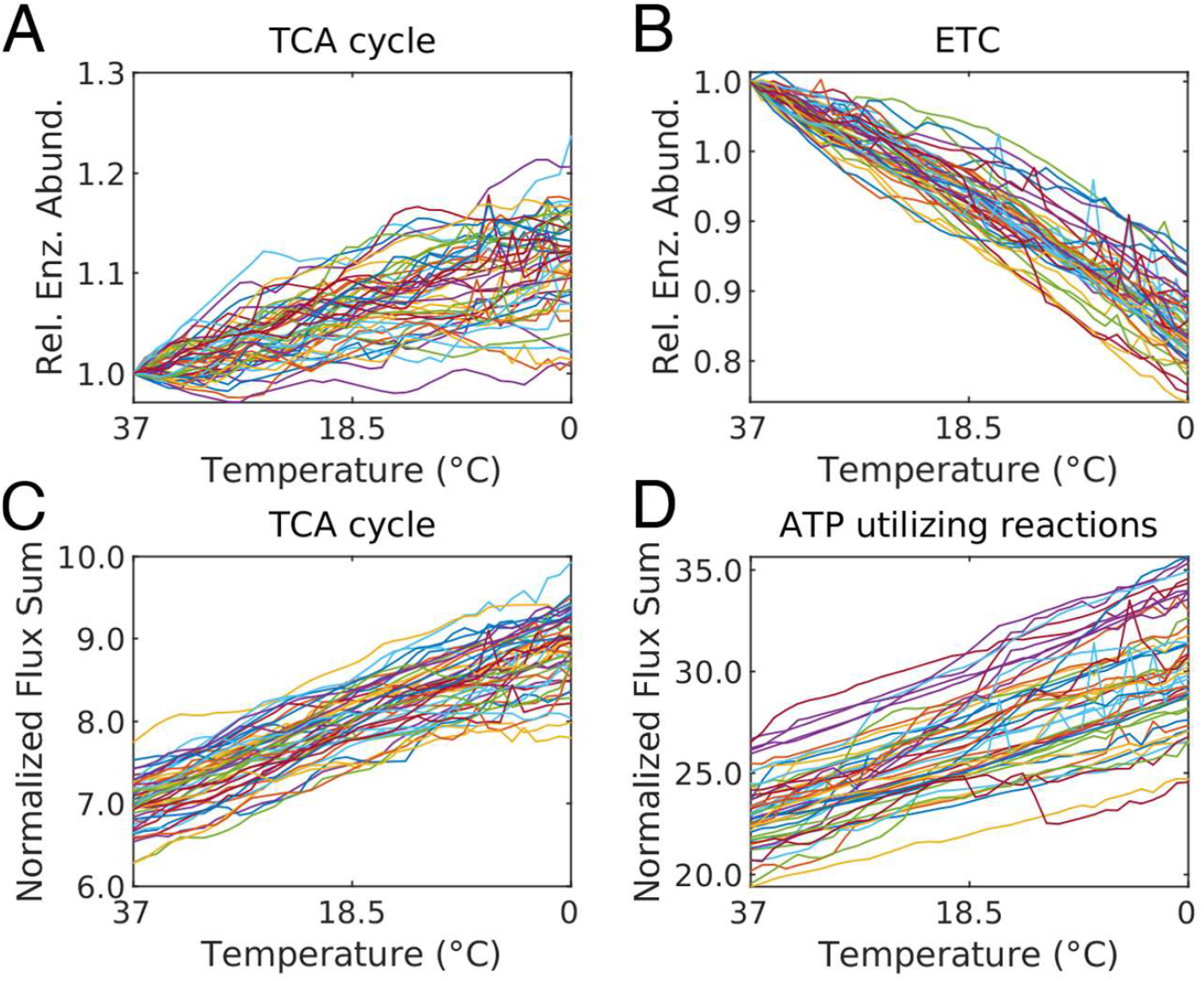
(A, B) Changes in the summed abundances of TCA cycle enzymes (A) and ETC enzymes (B) at stable steady states across temperatures. Each line represents one *x*_*ref*_. (C, D) Changes in the summed absolute values of TCA cycle reaction rates (C) and ATP-utilizing reaction rates (D), both normalized to glucose influx, at stable steady states across temperatures. Each line represents one *x*_*ref*_*ss*.

Because rate constants also vary with temperature, changes in enzyme abundance do not translate directly into proportional changes in flux. Therefore, to understand how enzyme adjustments maintain a high ATP yield during cooling, it is essential to examine the resulting metabolic fluxes rather than enzyme levels alone. We analyzed reaction rates normalized by glucose influx to investigate flux-level modulations arising from enzyme redistribution. The analysis revealed that, as temperature decreased, normalized flux through the TCA cycle increased, along with fluxes of reactions involved in ATP synthesis or consumption. Figures 5C and 5D show the summed reaction fluxes of the TCA cycle and ATP-utilizing reactions at stable steady states for each temperature. These findings indicate that the adjusted enzyme abundances enhanced ATP–ADP turnover, analogous to the rapid ATP–ADP exchange artificially imposed in the previous section. Together, these results suggest that maintaining a high ATP/ADP ratio through increased cofactor turnover allows cells to sustain high ATP yield even at low temperatures.

## Discussion

In this study, we computationally investigated how perturbations in environmental parameters influence the stability of a metabolic state. By varying rate constants in a temperature-dependent manner, we found that lowering the temperature caused a bifurcation and a steep drop in the ATP yield. This finding provides a model-based illustration of how complex metabolic systems can be destabilized by parameter perturbations (Chakrabarti et al., 2013; Murayama et al., 2017; Steuer et al., 2006).

One way to maintain a high ATP yield was to keep the ATP/ADP ratio constant through external control. Several studies have reported that coenzymes such as ATP and ADP exert a major influence on the perturbation responses of kinetic models (Hatakeyama & Furusawa, 2017; Himeoka & Mitarai 2022; Himeoka & Furusawa, 2025). In our model, fixing this cofactor ratio preserved a stable metabolic state across the temperature range and maintained a high ATP yield as shown in Figure 3C. We also demonstrated that the ATP yield can be maintained at a high level by adjusting enzyme abundances. Under such enzyme-tuning scenarios, the relative flux through ATP-utilizing reactions increased, keeping the ATP/ADP ratio nearly constant. Thus, regulation of enzyme abundances can, in principle, achieve the same effect as cofactor-ratio control.

Analysis of enzyme abundances to maintain the ATP yield revealed an increase in the TCA cycle enzymes and a decrease in ETC enzymes. This trend aligns with experimental reports that *E. coli* increases the TCA cycle enzyme levels and decreases ATP synthase subunits under cold temperature (Knapp et al., 2024; Phadtare & Inouye, 2004). This suggests that cells may modulate enzyme abundances *in vivo* to maintain the ATP/ADP ratio and prevent an abrupt shift in their metabolic state. Nevertheless, these findings require experimental validation, such as comprehensive fluxomic analyses that cover pathways beyond our model.

Our analysis perturbed only temperature, starting from parameters fitted to the 37 °C metabolic state. Therefore, the conclusions depend on that particular parameter set. Additionally, the model is limited to central metabolism and omits lipid pathways and many amino-acid biosynthesis routes that are reported to be important for cold adaptation (Phadtare & Inouye, 2008; Zhang & Gross, 2021). Network structure modifications in kinetic models are known to exert strong influences on responses for parameter perturbations (Hishida et al., 2023). Future work can probe cellular responses to cooling with higher quantitative accuracy using a more detailed network and more precise temperature dependence of parameters.

The same computational framework can be applied to environmental perturbations beyond temperature. By allowing kinetic parameters to vary with factors such as pH, osmotic pressure, or nutrient availability, we can reveal which metabolic functions become unstable and what operations— such as cofactor balance control or targeted enzyme tuning—can restore stability.

Overall, our results highlight the latent risk of destabilization in complex metabolic systems under environmental perturbations and identify cofactor balance—specifically the ATP/ADP ratio— as a critical variable for maintaining stability. Applying this approach to other external parameters may allow systematic identification of key variables that cells must regulate to preserve metabolic stability. Such insights would advance our understanding of the principles underlying biological robustness.

**Supplementary Figure S1.**
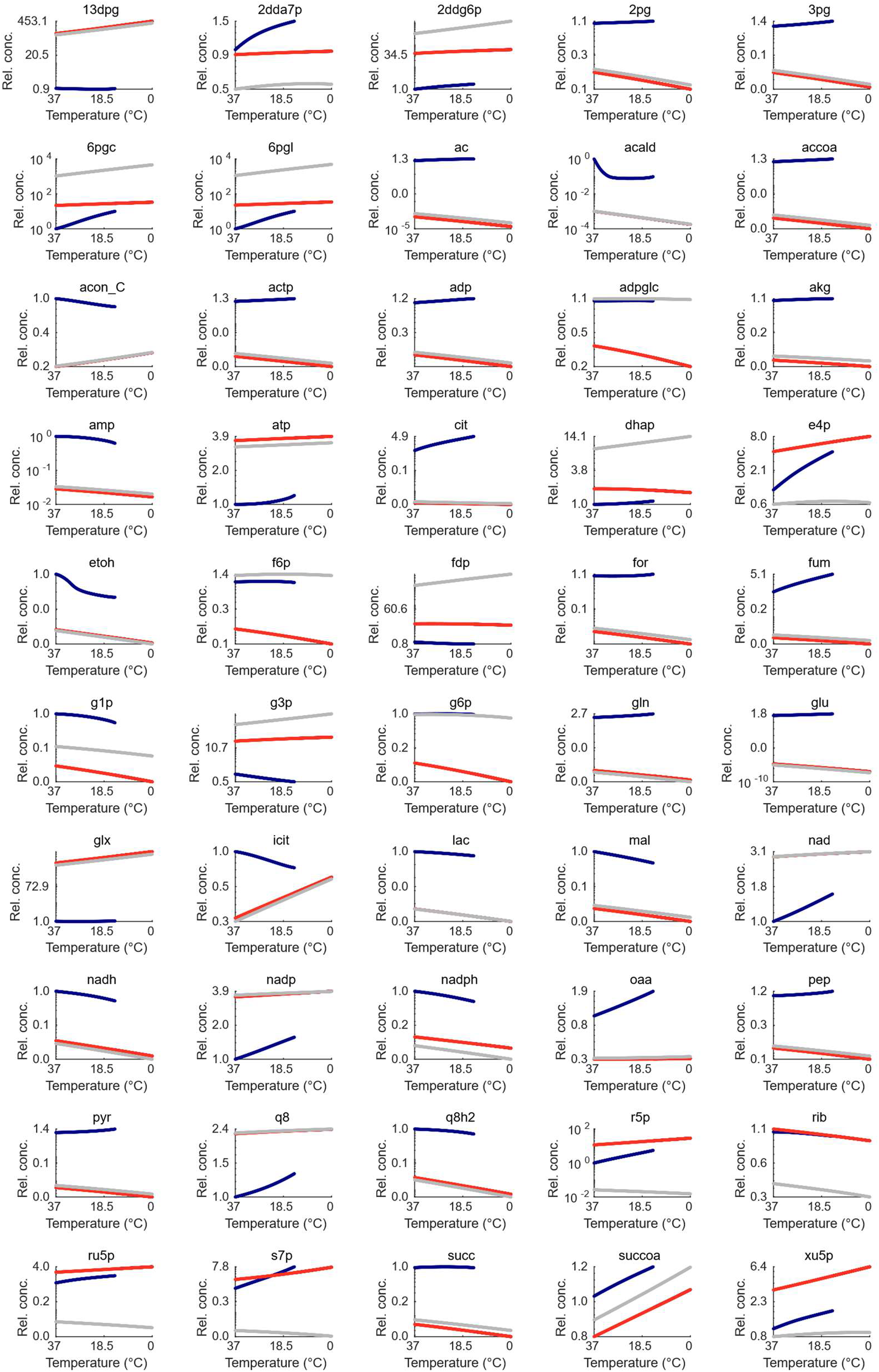
Bifurcation diagram of small chemicals. The blue line represents the steady state used for parameter fitting in Khodayari et al. (2014) at 37 °C, while the red and grey lines correspond to steady states that exist below 14.2 °C. The steady state at the blue line shifts to that represented by the red line due to the temperature-induced bifurcation.

**Supplementary Table S1.**
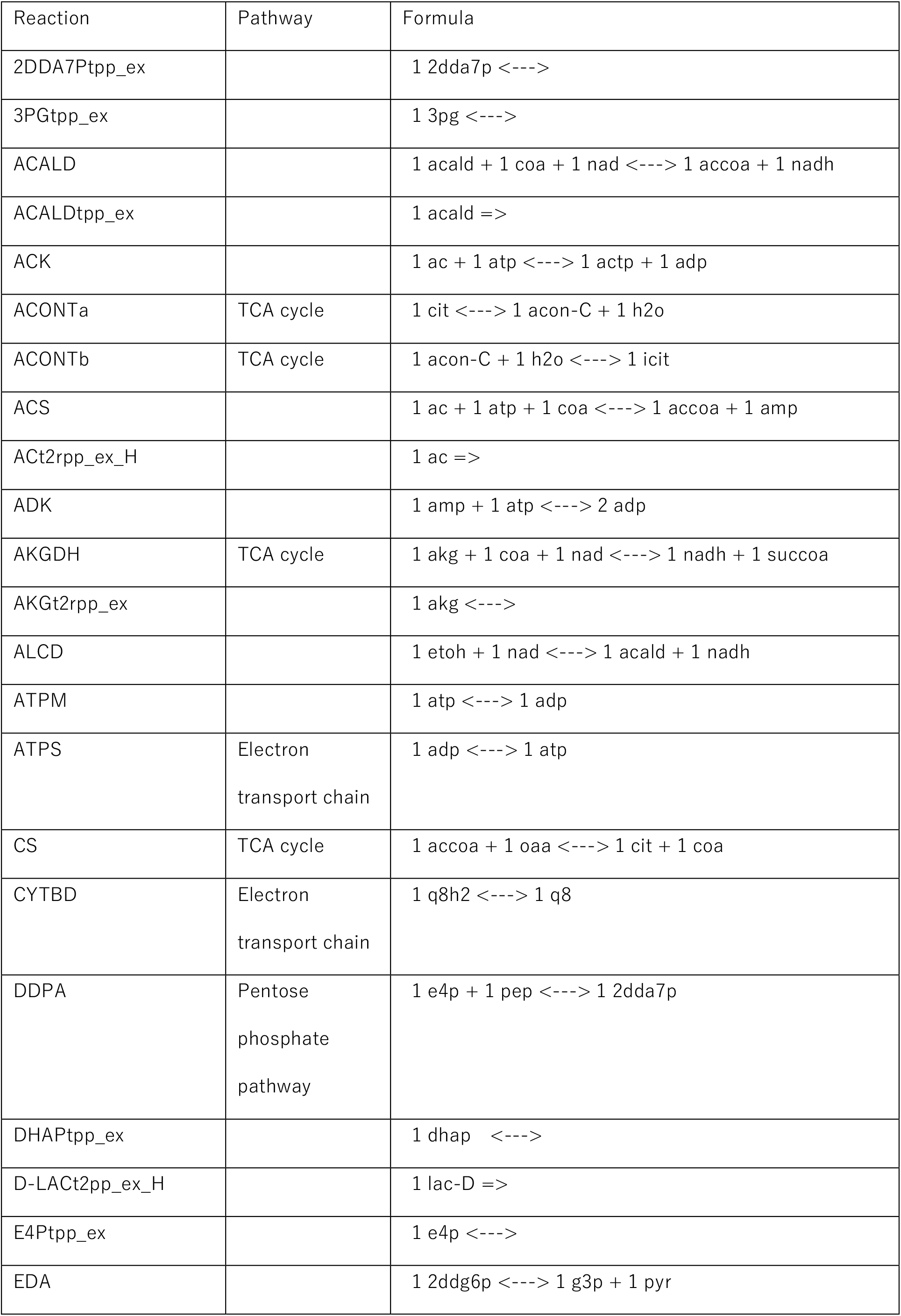
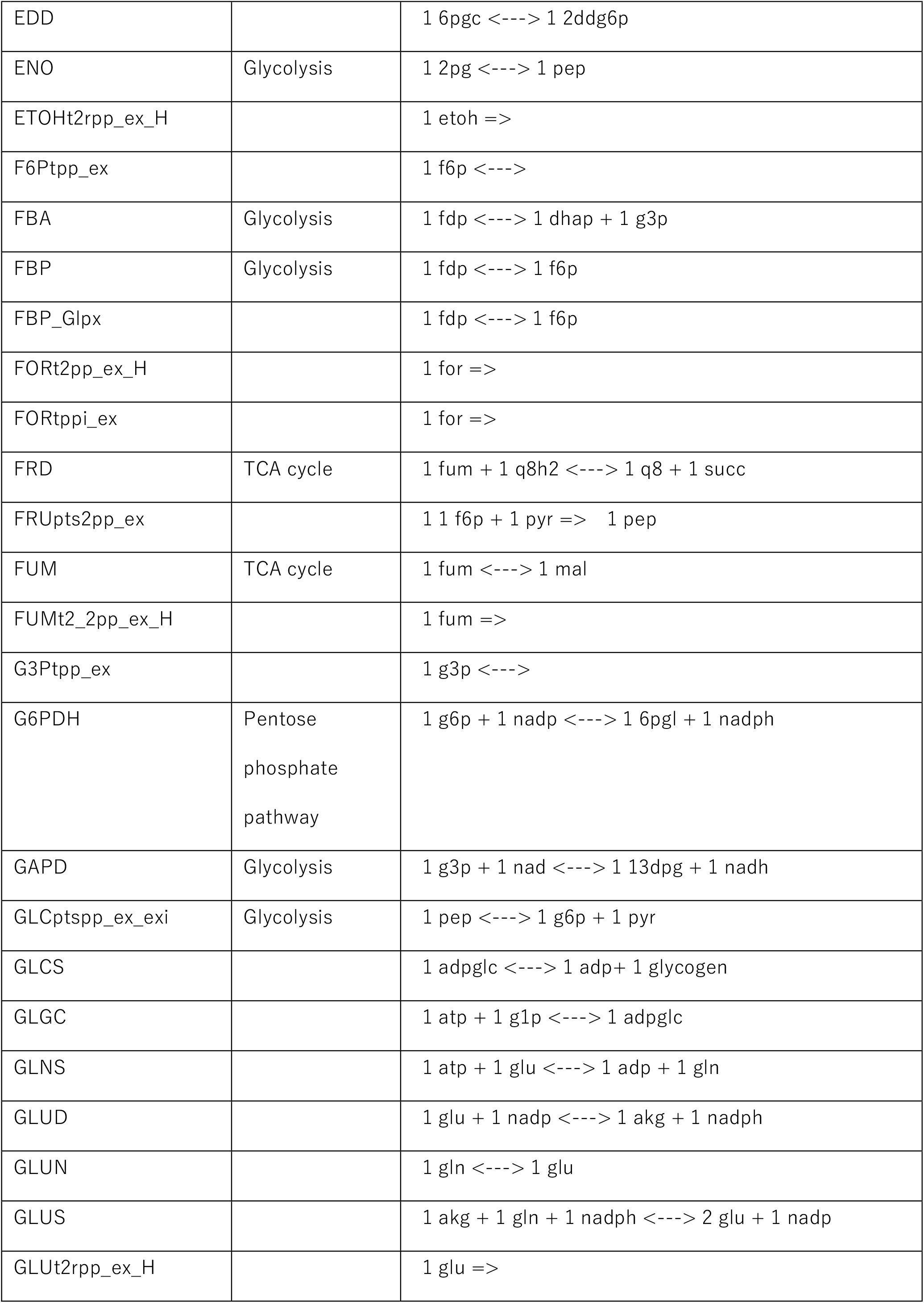
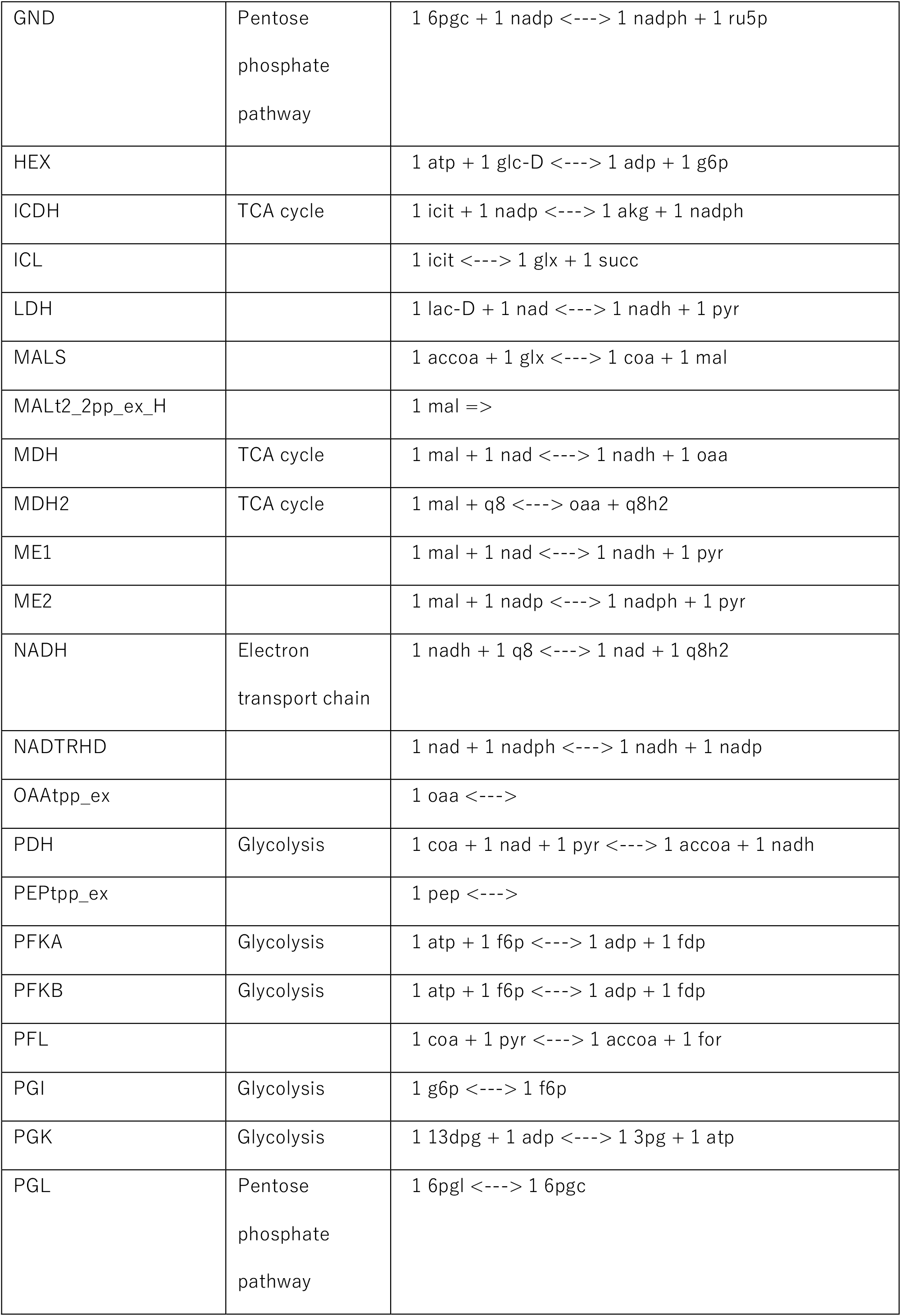
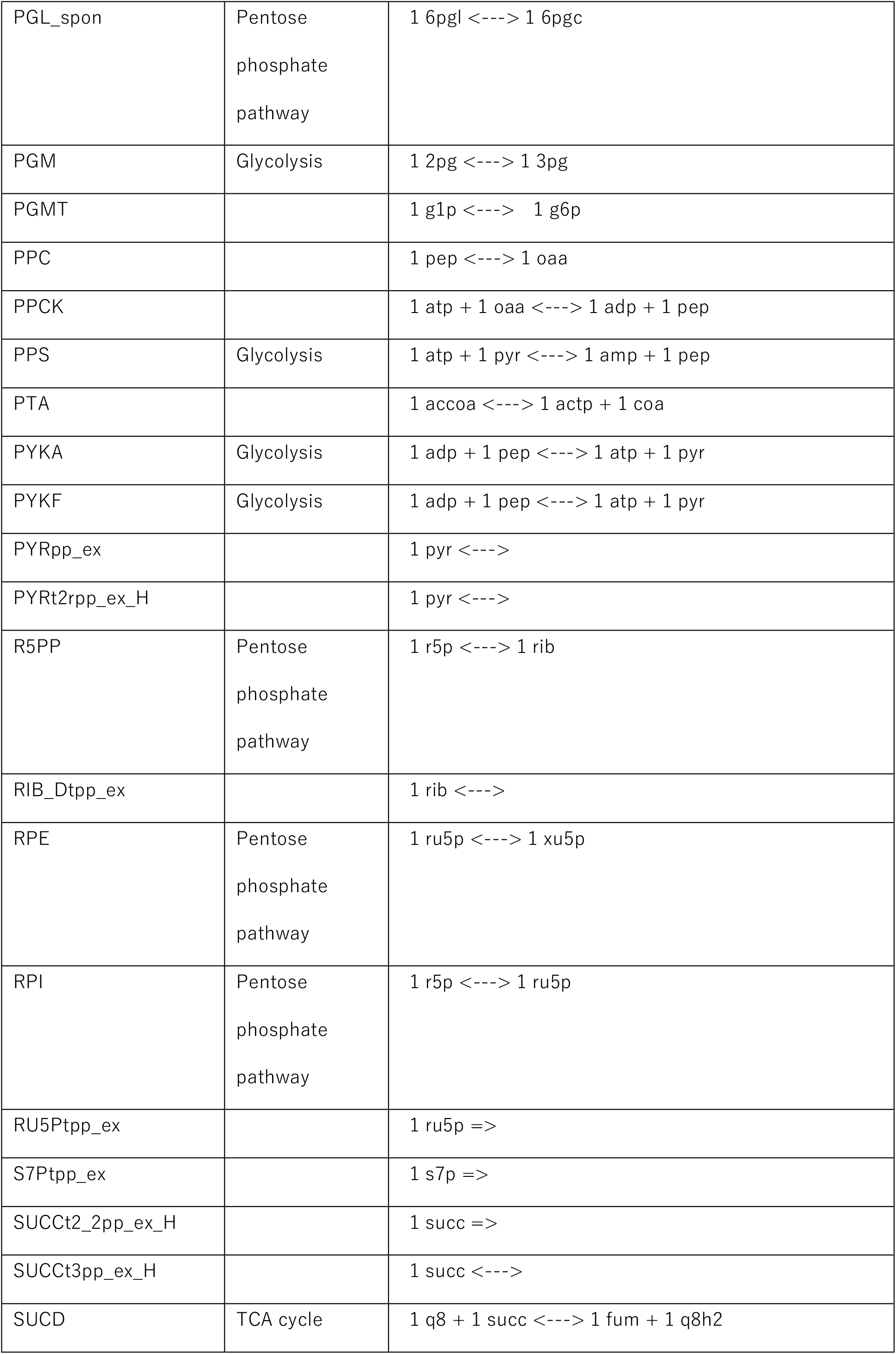
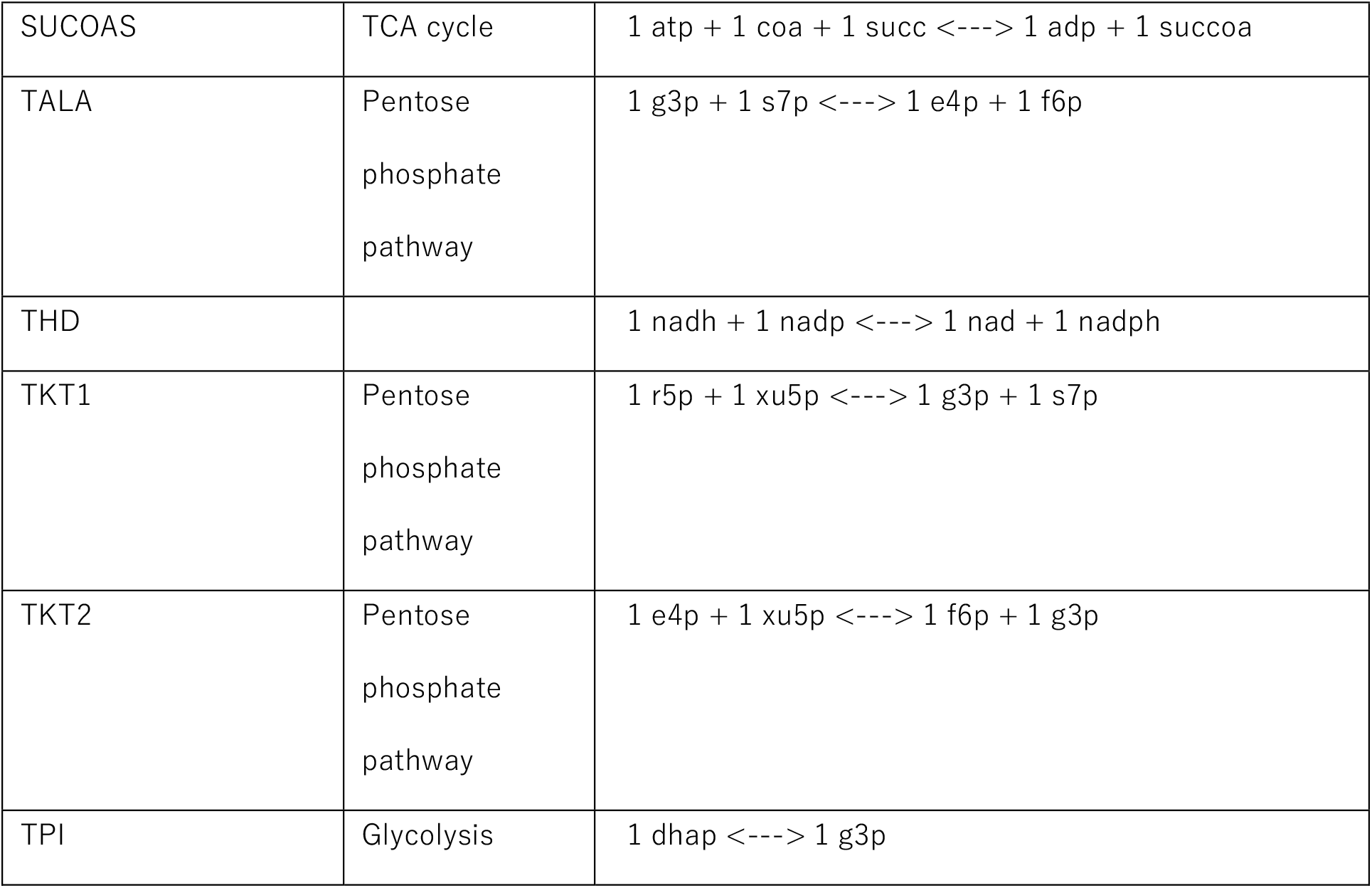
List of all reactions in the model used in our simulations.

**Supplementary Table S2.** List of all regulations in the model used in our simulations.

| Reaction | Regulator | Regulation Type |
| --- | --- | --- |
| ACALD | amp | Inhibition |
| EDA | 6pgc | Inhibition |
| EDA | g3p | Inhibition |
| FBA | dhap | Inhibition |
| FBP | adp | Inhibition |
| FBP | amp | Inhibition |
| FBP | pi | Inhibition |
| FBP | pep | Inhibition |
| FUM | cit | Inhibition |
| G6PDH | nadh | Inhibition |
| G6PDH | nadph | Inhibition |
| GLGC | amp | Inhibition |
| GLUN | adp | Inhibition |
| GLUN | atp | Inhibition |
| GLUN | glu | Inhibition |
| GLUN | nad | Inhibition |
| GLUN | nadh | Inhibition |
| GLUN | nadp | Inhibition |
| GLUN | nadph | Inhibition |
| GLUN | pi | Inhibition |
| GLUS | glu | Inhibition |
| GND | atp | Inhibition |
| GND | ru5p | Inhibition |
| GND | fdp | Inhibition |
| GND | nadp | Inhibition |
| ICDH | glx | Inhibition |
| ICDH | oxa | Inhibition |
| ICDH | pep | Inhibition |
| ICL | akg | Inhibition |
| ICL | acon | Inhibition |
| ICL | pep | Inhibition |
| ICL | succ | Inhibition |
| LDH | atp | Inhibition |
| PGI | 6pgc | Inhibition |
| PGI | pep | Inhibition |
| PGMT | accoa | Inhibition |
| PGMT | g1p | Inhibition |
| PPC | mal | Inhibition |
| PPC | pep | Inhibition |
| PPCK | atp | Inhibition |
| PPCK | pep | Inhibition |
| PPS | akg | Inhibition |
| PPS | adp | Inhibition |
| PPS | amp | Inhibition |
| PPS | oaa | Inhibition |
| PPS | pep | Inhibition |
| PTA | adp | Inhibition |
| PTA | atp | Inhibition |
| PTA | nadh | Inhibition |
| PTA | nadph | Inhibition |
| PTA | pep | Inhibition |
| PTA | pyr | Inhibition |
| PYKA PYKF | atp | Inhibition |
| PYKA PYKF | succoa | Inhibition |
| RPI | amp | Inhibition |
| TALA | pi | Inhibition |

## Materials and Methods

### Model Description

The model used in this analysis is based on the model from Himeoka et al. (2025), which is a modified version of the Khodayari et al. (2014) model, adjusted to ensure the existence of a stable steady state. In the present study, the parameters of inflow and outflow reactions were further modified so that glucose serves as the sole carbon source in the temperature range used for the simulations. The model includes glycolysis, the TCA cycle, the pentose phosphate pathway (PPP), glutamine synthesis, the electron transport chain (ETC), and the Entner-Doudoroff pathway. See Supplementary Table S1 and S2 for complete reaction formulae, pathway components, and regulations.

In our model, all reactions are decomposed into elementary reactions that represent substrate-enzyme binding and dissociation. This is illustrated using a simple reversible reaction as follows:

For a reversible reaction A + B ⇌ C catalyzed by an enzyme E, we write:

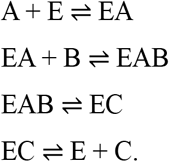

All steps are reversible, and the forward and reverse directions are treated as distinct elementary reactions. The total concentration of free and complex enzymes does not change in time:

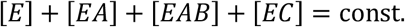

The rates of these elementary reactions are calculated based on mass-action kinetics. Note that in our model, chemical concentrations are dimensionless because they are normalized to 1.0 at the reference steady state as a standard procedure in ensemble modeling (Khodayari et al., 2014; Tran et al., 2008).

In our model, the time evolution of concentrations is described by an ODE system

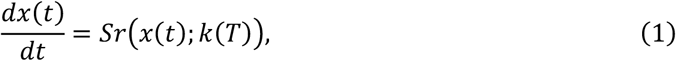

where *x* is the concentration vector of small molecules, free enzymes, and substrate-enzyme complexes; *S* is the stoichiometric matrix; *k*(*T*) is the set of rate constants at temperature *T*; and *r*(*x*) is the vector of reaction rate functions. All reactions are decomposed into elementary reactions, and each reaction rate is modeled by the law of mass action

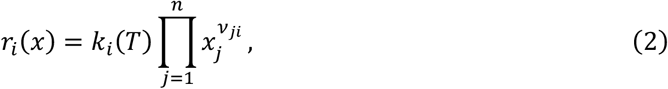

with exponents *v*_*ji*_ ∈ {0,1} determined by the stoichiometry and *k*_*i*_(*T*) the temperature-dependent rate constants (specified in the next section).

ODE simulations were performed using the ode15s function in MATLAB (The MathWorks Inc., 2023).

### Temperature-Dependent Rate Constants

Rate constants were adjusted with the Eyring-Polanyi equation

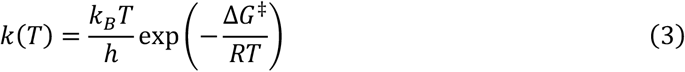

for each temperature. Here, *k* is the rate constant, *h* is the Planck constant, *k*_*B*_ is the Boltzmann constant, *R* is the gas constant, and *T* is the absolute temperature (*K*). The activation free energy Δ*G*^‡^ for each reaction is back-calculated from parameter values obtained in Khodayari et al. 2014 using the following equation:

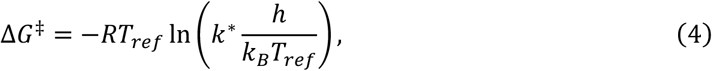

with *T*_*ref*_ = 310.15, and *k*^∗^ is the rate constant at *T*_*ref*_. Based on this value, reaction rate constants at altered temperatures were calculated.

### Maintaining the ATP/ADP Ratio

To forcibly maintain the ATP/ADP ratio while lowering the temperature, we added a new reversible reaction that exchanges ATP and ADP to the model. The rate constant for this exchange was set to 10^6^, a value larger than the median of all rate constants at 37 °C. During low-temperature simulations, this constant was not adjusted for temperature dependence. The concentrations of the free metabolites at the steady state are normalized to unity in the ensemble modeling procedure carried out for the model construction in Khodayari et al. (2014). Thus, ATP/ADP ratio in the original model is 1:1 in the scale of normalized concentrations. Therefore, the force-balancing ATP/ADP ratio to unity corresponds to the recovery of the ATP/ADP ratio in the original model.

### Perturbed Initial Condition Sampling Procedure

To identify multiple stable steady states in the system, we perform numerical integrations starting from perturbed initial conditions with specified temperature parameters. These initial conditions are generated by multiplying the concentrations of an original steady state by random values with ensuring that the conserved quantities are not changed.

If a vector *d* satisfies

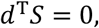

then the inner product of *d* and a concentration vector *x* does not change in time, since

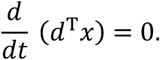

If random perturbations are applied directly to the concentrations of the original steady state *x*_*org*_, the resulting concentration vector *x*_*rand*_ will generally have conserved quantities different from those of *x*_*org*_. To obtain perturbed concentrations that preserve the conserved quantities of *x*_*org*_, we search for the point closest to *x*_*rand*_ that satisfies the conservation constraints.

This is formulated as the following optimization problem:

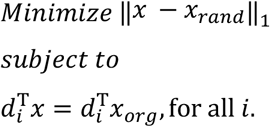

Where {*d*_*i*_} form a basis of ker *S*^*T*^. This optimization is done using Gurobi Optimizer (Gurobi Optimization, 2024).

### Steady State Sampling Procedure

To identify enzyme abundances that yield a stable metabolic state with low-temperature parameters, we sampled steady states for each temperature-specific parameter set and computed the corresponding enzyme abundances. Each reaction is decomposed into elementary reactions whose reactants are (i) substrate and enzyme, (ii) substrate and substrate-enzyme complex, or (iii) substrate-enzyme complex only. Therefore, each reaction rate function is a first- or second-order monomial in its variables.

Fixing concentrations of substrates (small molecules) at their values near the original steady state turns every rate into a linear function of the free-enzyme or substrate-enzyme complex concentrations. *x*_*enzyme*_ is a vector of the remaining variables, i.e., the concentrations of free enzymes and substrate-enzyme complexes. Using a matrix *V*, whose non-zero values are determined by the temperature-specific rate constants and the fixed substrate concentrations, we have

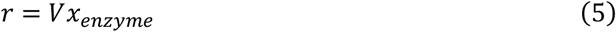

where *r* is the vector of reaction rates.

For example, if the system has only one reaction A + B ⇌ C catalyzed by an enzyme E, reaction rate functions are

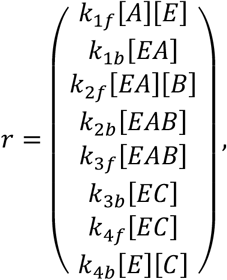

where *k*_i_ is rate constant of forward or backward direction of each elementary reaction. When the substrate concentrations are fixed at ([*A*]^∗^, [*B*]^∗^, [*C*]^∗^)^T^, the rate functions can be rewritten as

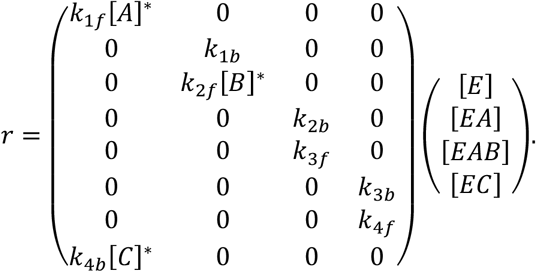

Using submatrix of the stoichiometric matrix corresponding to the remaining variables, denoted as *S*_*enzyme*_, the time-course change of *x*_*enzyme*_ can be written as

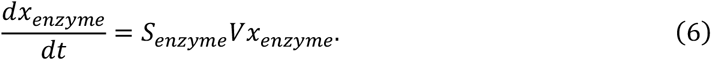

Therefore, steady-state solutions are obtained just by solving

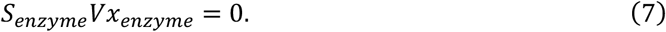

Eq. (7) is less-determined linear equation when *S*_*enzyme*_*V* is not full-rank. In that case, Eq. (7) has infinitely many solutions. Among such solutions of Eq. (7), we select the one closest to a reference point (*x*_*ref*_), which is randomly sampled around the enzyme abundances of the original steady state. We also require steady states that

1. Glucose is a single energy resource, and transport reactions of other chemicals work as efflux.
2. The total enzyme abundance of all reactions is equal to that of the original steady state.

Steady states satisfying those conditions are obtained by solving the following optimization problem:

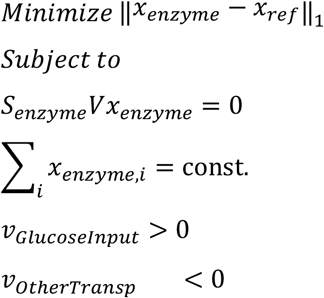

The solution *x*_*enzyme*_ obtained from this optimization, together with the fixed small molecule concentrations near the 37 °C steady state, yields a steady state of the full system. These optimizations are done using Gurobi Optimizer (Gurobi Optimization, 2024).

### Stability Assessment of Steady States

The stability of each steady state found by the optimization was determined by computing the eigenvalues of the system’s Jacobi matrix at the steady state. The model contains 98 conserved quantities, so the full 643 × 643 Jacobi matrix has 98 zero eigenvalues. The zero eigenvalues are often tricky to distinguish from the very small, non-zero eigenvalues due to numerical errors. Thus, we avoided obtaining zero eigenvalues by pre-solving the dependent variables using the conserved quantities. Note that one conserved quantity leads to a single linear equation. For each conserved quantity, we choose a single chemical species and solve for its concentration as a function of the remaining species’ concentrations. From this pre-solved ODE system, we obtain the reduced 545 × 545 Jacobi matrix. A steady state was classified as unstable if the largest real part of any eigenvalue of the reduced Jacobi matrix was positive. All eigenvalues were computed with interval-arithmetic routines in INTLAB, ensuring that numerical errors did not flip the sign of any eigenvalue (Rump, 1999).

## Acknowledgements

This work is supported in part by JSPS KAKENHI (23KJ1324 to A. H.; 22K15069, 24H01118 and 25H01390 to Y. H.; 22K21344 and 23H0247 to C. F.), and GteX Program Japan Grant No. JPMJGX23B4.

## Notes

### Competing Interest Statement

The authors have declared no competing interest.

